# Nitrogen-fixing populations of Planctomycetes and Proteobacteria are abundant in the surface ocean

**DOI:** 10.1101/129791

**Authors:** Tom O. Delmont, Christopher Quince, Alon Shaiber, Özcan C. Esen, Sonny TM Lee, Sebastian Lücker, A. Murat Eren

## Abstract

Nitrogen fixation in the surface ocean impacts the global climate by regulating the microbial primary productivity and the sequestration of carbon through the biological pump. Cyanobacterial populations have long been thought to represent the main suppliers of the bio-available nitrogen in this habitat. However, recent molecular surveys of nitrogenase reductase gene revealed the existence of rare non-cyanobacterial populations that can also fix nitrogen. Here, we characterize for the first time the genomic content of some of these heterotrophic bacterial diazotrophs (HBDs) inhabiting the open surface ocean waters. They represent new lineages within Planctomycetes and Proteobacteria, a phylum never linked to nitrogen fixation prior to this study. HBDs were surprisingly abundant in the Pacific Ocean and the Atlantic Ocean northwest, conflicting with decades of PCR surveys. The abundance and widespread occurrence of non-cyanobacterial diazotrophs in the surface ocean emphasizes the need to re-evaluate their role in the nitrogen cycle and primary productivity.

## Introduction

Marine microbial communities play a critical role in biogeochemical fluxes and climate [1, 2, 3], but their activity in the euphotic zone of low latitude oceans is often limited by nitrogen availability [4, 5]. Thus, biological fixation of gaseous dinitrogen in the surface ocean is a globally important process that contributes to the ocean’s productivity and enhances the sequestration of carbon through the biological pump [6, 7]. Microbial populations that can fix nitrogen are termed diazotrophs, and although they encompass a wide range of archaeal and bacterial lineages [8, 9], diazotrophs within the bacterial phylum Cyanobacteria in particular are considered to be responsible for a substantial portion of nitrogen input in the surface ocean [10, 11, 12]. Studies employing cultivation and flow cytometry [13, 14, 15, 16, 17] characterized multiple cyanobacterial diazotrophs and shed light on their functional life-styles [18, 19, 20]. PCR surveys of the nitrogenase reductase gene *nifH* have indicated that the ability to fix nitrogen is also found in lineages of the phyla Proteobacteria, Firmicutes, and Spirochetes [9, 21, 22], suggesting the presence of heterotrophic bacterial diazotrophs (HBDs) that contribute to the introduction of nitrogen in the surface ocean. Quantitative surveys of non-cyanobacterial *nifH* genes indicated that HBDs are diverse but relatively rare in open ocean [23, 24, 25, 26, 27, 28], and efforts to access their genomic contents through cultivation have not yet been successful, limiting our understanding of their ecophysiology.

Here, we used metagenomic co-assembly and binning strategies to create a non-redundant database of archaeal, bacterial and eukaryotic genomes from the TARA Oceans project [29]. We characterized nearly one thousand microbial genomes from the surface samples of four oceans and two seas, which revealed novel nitrogen-fixing Planctomycetes and Proteobacteria populations. In contrast to previous findings of PCR-dependent surveys, our analyses indicate that putative HBDs are not only diverse but also abundant in the open surface ocean habitat.

## Results

The 93 TARA Oceans metagenomes we analyzed represent the planktonic size fraction (0.2-3 micrometer) of 61 surface samples and 32 samples from the deep chlorophyll maximum layer of the water column (Table S1). 30.9 billion metagenomic reads (out of 33.7 billion) passed the quality control criteria. We performed 12 metagenomic co-assemblies (1.14-5.33 billion reads per set) using geographically bounded samples (Figure 1), and identified 42.2 million genes in scaffolds longer than 1,000 nucleotides (Table S2 summarizes the assembly statistics). We applied a combination of automatic and manual binning to each co-assembly output, which resulted in 957 manually curated, non-redundant metagenome-assembled genomes (MAGs) (Figure 1).

**Figure 1.**
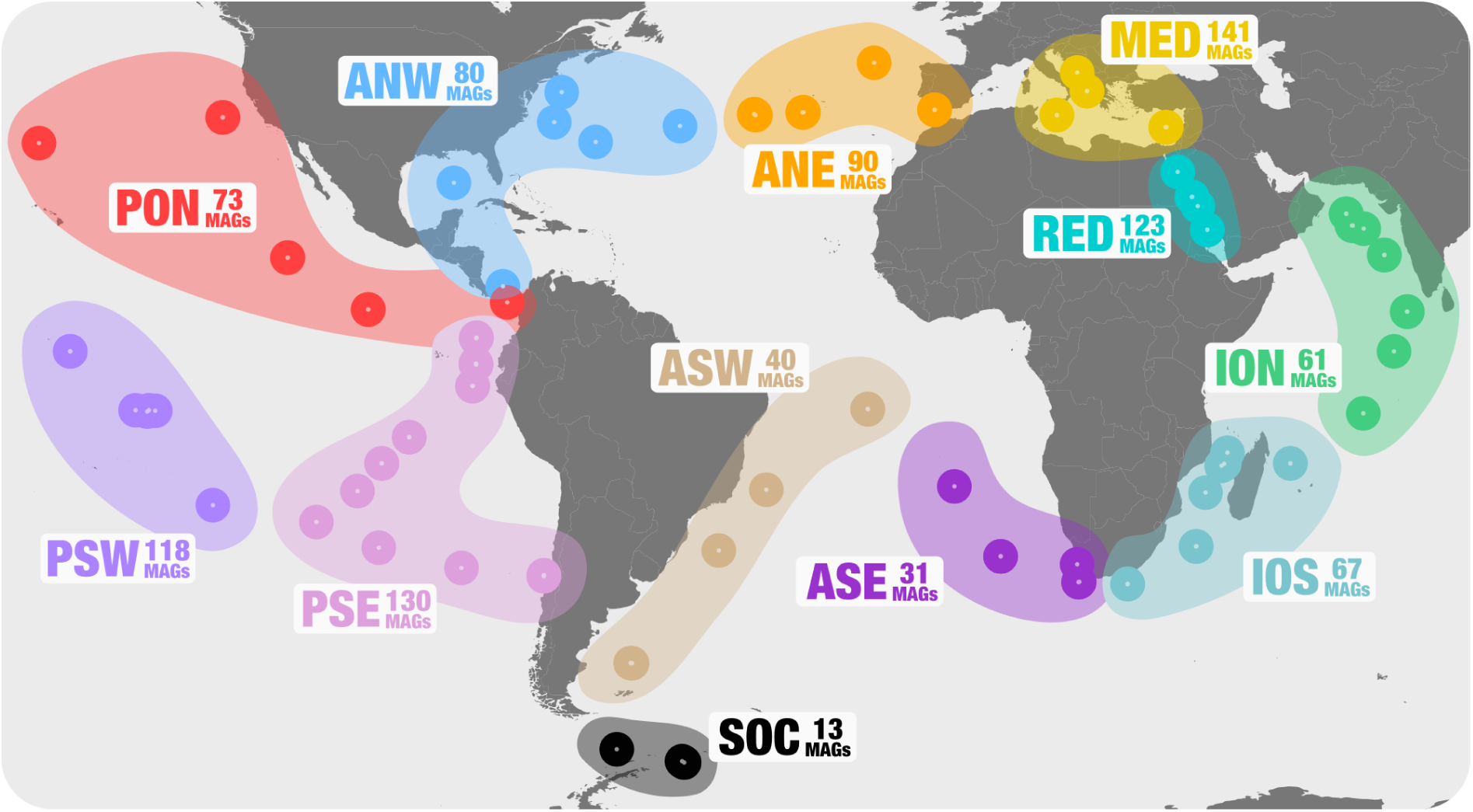
Geographically bounded metagenomic co-assemblies. Dots in the map correspond to the geographic location of 93 metagenomes from the TARA Oceans project. Each dot is associated with a metagenomic set (corresponding to a geographic region) for which we performed a metagenomic co-assembly (n=12). The number of metagenome-assembled genomes (MAGs) we characterized from each metagenomic set varied from 13 to 141 after the removal of redundant MAGs, for a total of 957 non-redundant MAGs encompassing the three domains of life.

Our MAGs belonged to the domains Bacteria (n=820), Eukaryota (n=72) and Archaea (n=65) (Table S3), and recruited 2.11 billion raw reads (6.84% of the dataset) when we mapped metagenomic data back to this collection. The genomic completion and redundancy estimates for archaeal and bacterial MAGs based on domain specific single-copy core genes averaged to 79% and 76.1%, respectively, and resolved to phyla Proteobacteria (n=432), Bacteroidetes (n=113), Euryarchaeota (n=65), Verrucomicrobia (n=65), Planctomycetes (n=43), Actinobacteria (n=37), Chloroflexi (n=34), Candidatus Tarascapha (n=27), Acidobacteria (n=6), Cyanobacteria (n=6), Spirochaetes (n=5), Firmicutes (n=2), Ignavibacteriae (n=1), and diverse members of the Candidate Phyla Radiation (n=4). We could assign only 6.33% of the bacterial and archaeal MAGs to described genera. Eukaryotic MAGs were substantially larger than bacterial and archaeal MAGs (7.24 Mbp versus 2.26 Mbp and 1.47 Mbp, respectively) and were dominated by a small number of genera: *Micromonas* (n=14), *Emilliana* (n=14), *Bathycoccus* (n=8) and *Ostreococcus* (n=4). Recovery of these MAGs complements decades of cultivation efforts by providing genomic context for lineages missing in culture collections (e.g., Euryarchaeota, Candidatus Tarascapha, and the Candidate Phyla Radiation), and allowed us to search for diazotrophs within a large pool of mostly novel marine microbial populations.

### Genomic stability of a well-studied nitrogen-fixing symbiotic population at large scale

Our genomic collection included six cyanobacterial MAGs, one of which (ASW 00003), contained genes that encode the catalytic (*nifH*, *nifD*, *nifK*) and the biosynthetic proteins (*nifE*, *nifN* and *nifB*) required for nitrogen fixation [8]. ASW 00003, which we recovered from the Atlantic southwest metagenomic co-assembly, showed remarkable similarity to the genome of the symbiotic cyanobacterium Candidatus *Atelocyanobacterium thalassa* [30] isolated from the North Pacific gyre (GenBank accession: CP001842.1). Besides their comparable size of 1.43 Mbp (ASW 00003) and 1.46 Mbp (isolate), their average nucleotide identity was 99.96% over the 1.43 Mbp alignment. Ca. *A. thalassa* is a diazotrophic taxon that lacks key metabolic pathways and lives in symbiosis with photosynthetic eukaryotic cells [19, 31]. While the high genomic similarity between ASW 00003 and the Ca. A. thalassa isolate genome demonstrates the accuracy of our metagenomic workflow, this finding also emphasizes the genomic stability of a well-studied nitrogen-fixing symbiotic population at large scale, possibly due to a strong selective pressure from the eukaryotic hosts.

### Genomic evidence for nitrogen fixation by Planctomycetes and Proteobacteria

Besides the cyanobacterial MAG, we also identified seven Proteobacteria and two Planctomycetes MAGs in our collection that contained the complete set of genes for nitrogen fixation. To the best of our knowledge, these MAGs (HBD-01 to HBD-09) represent the first genomic evidence of putative HBDs inhabiting the surface of the open ocean (Figure 2, Panel A). They were obtained from the Pacific Ocean (n=6), Atlantic Ocean (n=2), and Indian Ocean (n=1), and displayed relatively large genome sizes (up to 6 Mbp and 5,390 genes) and a GC-content ranging from 50% to 58.7%. One of the Proteobacterial MAGs resolved to the genus *Desulfovibrio* (HBD-01). The remaining MAGs from this phylum represented new lineages within the orders Desulfobacterales (HBD-02), Oceanospirillales (HBD-03; HBD-04; HBD-05) and Pseudomonadales (HBD-06; HBD-07) (Figure 2, Panel A). The phylogenetic assignment of one Planctomycetes MAG (HBD-08) with a low completion estimate (33.5%) could not be resolved beyond the phylum level, possibly due to missing phylogenetic marker genes for taxonomic inferences. However, the length of this MAG (4.03 Mbp) suggests that its completion based on single-copy core genes may have been underestimated, as we have observed in previous studies [32, 33]. The closest match for the second Planctomycetes MAG (HBD-09) was the family Planctomycetaceae (order Planctomycetales) when using a collection of single copy genes. This MAG contained a large fragment of the 16S rRNA gene (1,188 nt; available in Table 4) for which the best match to any characterized bacterium in the NCBI’s non-redundant database was *Algisphaera agarilytica* (strain 06SJR6-2, NR 125472) with 88% identity. The novelty of the 16S rRNA gene sequence suggests that this MAG represents a new order (and possibly a new class) within the Planctomycetes.

**Figure 2.**
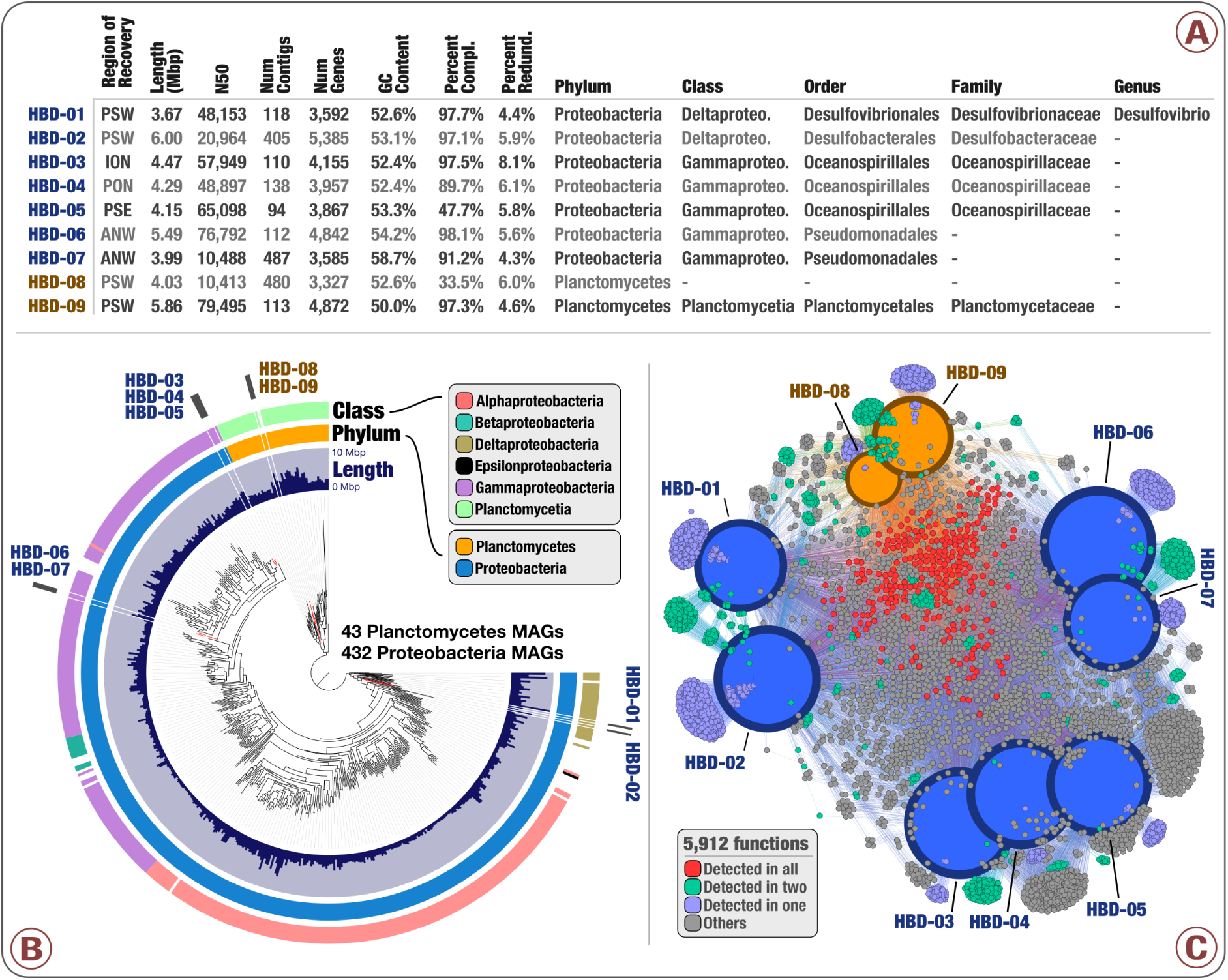
Nexus between phylogeny and function of HBDs. (A) genomic feature summary of the nine HBDs. (B) Phylogenomic analysis of 432 Proteobacteria MAGs and 43 Planctomycetes MAGs in the non-redundant genomic database (including the nine HBDs) using a collection of 37 phylogenetic marker gene families. Layers surrounding the phylogenomic tree indicate genome size and taxonomy of each MAG at the phylum and class level. (C) Functional network of the nine HBDs base upon a total of 5,912 identified gene functions. Size and colour of genomic nodes represent the number of detected functions and MAG taxonomy, respectively. Finally, the colour of functional nodes indicates their occurrence in the different HBDs.

We placed the nine HBDs in a phylogenomic analysis of the 432 Proteobacteria and 43 Planctomycetes MAGs using a set of 37 phylogenetic marker genes (Figure 2, Panel B; interactive interface at https://anviserver.org/merenlab/tarahbds). The two Deltaproteobacteria HBDs were closely related to each other, but not adjacent in the phylogenomic tree. The HBDs within Oceanospirillales (n=3), Pseudomonadales (n=2) and Planctomycetes (n=2) formed three distinct phylogenomic lineages. These results suggest that closely related populations of diazotrophs inhabit the surface ocean, and nitrogen fixation genes occur sporadically among diverse putatively heterotrophic marine microbial lineages.

Our initial binning results included 120 redundant MAGs that we observed multiple times in independent co-assemblies (Table S5). Although they are not present in our final collection of 957 non-redundant MAGs (allowing for a better assessment of the relative abundance of microbial populations), we used this redundancy to investigate the stability of the phylogeny and functional potential of populations recovered from multiple geographical regions. For instance, we characterized the genomic content of HBD-06 from the Atlantic northwest (5.49 Mbp) and from each of the three Pacific Ocean regions (5.55 Mbp, 5.33 Mbp, and 5.29 Mbp in the regions PON, PSW, and PSE, respectively) (Table S5). Average nucleotide identity values between the Atlantic MAG and the three Pacific MAGs ranged from 99.89% to 99.97% over more than 97% of the genome length. We observed similar trends for HBD-07 and HBD-09 (Table S5). The complete set of nitrogen fixation genes was present in all these redundant MAGs, demonstrating a large-scale stability of this functional trait in these HBDs.

The proportion of genes of unknown function was on average 27.6(+/- 2.63)% for the Proteobacteria HBDs and 49.3(+/- 0.5)% for the Planctomycetes HBDs, exposing an important gap in our functional understanding of especially the latter taxonomic group of diazotrophs. The grand total of 37,582 genes we identified in the nine HBDs encoded 5,912 known functions (Table S6) and the network analysis of HBDs based on known functions organized them into four distinct groups corresponding to Deltaproteobacteria, Oceanospirillales, Pseudomonadales and Planctomycetes (Figure 2, panel C), mirroring the results of our phylogenomic analysis. A large number of functions we identified in these HBDs (4,224 out of 5,912) were unique to either of the four groups (Figure 2, Panel C; Table S6). The relatively weak overlap of known functions between these groups indicates that the ability to fix nitrogen in marine populations may not be associated with a tightly defined functional life style.

Functional differences between HBDs included multiple traits within the biological nitrogen cycle. For instance, hydrolysis of urea to ammonia and CO2 was characteristic to the Gammaproteobacteria HBDs, while oxidation of glycine to ammonia was only detected in the Deltaproteobacteria HBDs and the Planctomycetes HBD-08 (Table S6). These pathways might represent alternative ammonia sources to avoid the costly nitrogen fixation. In addition, we detected the complete denitrification pathway in the representative MAG and all three genomic replicates of HBD-06, suggesting that widespread HBDs may also be involved in nitrogen loss in the surface ocean. However, we only identified this pathway in HBD-06 and a non-diazotroph MAG (ION 00025 affiliated to the genus *Labrenzia*). Thus, the scarcity of nitrogen fixation (10/957) and denitrification (2/957) pathways in our genomic database suggest the metabolic potential of HBD-06 to be highly singular. Other functional differences between HBDs included genes related to the regulation of nitrogen fixation. Single copies of the coding genes nifX, nifQ, nifO, nifW and nifT were characteristic to Oceanospirillales and Pseudomonadales HBDs, while the two-component nitrogen fixation transcriptional regulator fixJ was only detected in the Planctomycetes HBDs. We also identified a relatively small set of 271 functions (4,608 genes) common to the nine HBDs (Figure 2, panel C). Besides various housekeeping genes and the full set of genes for nitrogen fixation, they included genes coding for chemotaxis and flagellar proteins, transporters for ammonia, phosphate and molybdenum (required for nitrogen fixation), and multiple nitrogen regulatory proteins (Table S6). They also included 477 genes coding for transcriptional regulators. HBDs also shared complete pathways for biosynthesis of acyl-CoA and acetyl-CoA (beta-oxidation and pyruvate oxidation), glycolysis, ABC-2 type transport systems, and a two-component regulatory system for phosphate starvation response (Table S5). Thus, the HBDs we characterized appeared to be involved in different steps of the nitrogen cycle and possessed distinct strategies regulating nitrogen fixation, but shared traits related to energy conservation, motility, nutrient acquisition and gene regulatory processes. Swimming motility was a common trait we observed in all the HBDs, which may be an indication of a free-living life style rather than the symbiotic lifestyles observed in some cyanobacterial diazotrophs.

### The taxonomy of HBDs is coherent with the phylogeny of nitrogen fixation genes

Our phylogenetic analysis of the catalytic genes *nifH* and *nifD* from a wide range of diazotrophs placed our HBDs in four distinct lineages (Figure 3). We also included in this analysis the genomic replicates we had removed from our non-redundant genomic collection. These replicates clustered with their representative MAGs in the phylogenetic tree, which indicates that widespread HBDs maintain near-identical nitrogen fixation genes. HBD-01 (Desulfovibrio) and HBD-02 (Desulfobacterales) were clustered with close taxonomic relatives. In addition, the Gammaproteobacteria HBDs were most closely related to reference genomes of the genera *Pseudomonas* and *Azotobacter* from the same class. The agreement between their taxonomy and placement in the functional gene-based phylogeny suggests that diazotrophic traits in the heterotrophic bacterial populations inhabiting the surface ocean may have been transferred vertically rather than horizontally. Finally, the *nifD* and *nifH* genes we identified in the Planctomycetes HBDs formed distinct phylogenetic clusters, which was particularly apparent for *nifD* (Figure 3, panel A). These results suggest nitrogen fixation represents a functional trait rooted within the phylum Planctomycetes.

**Figure 3.**
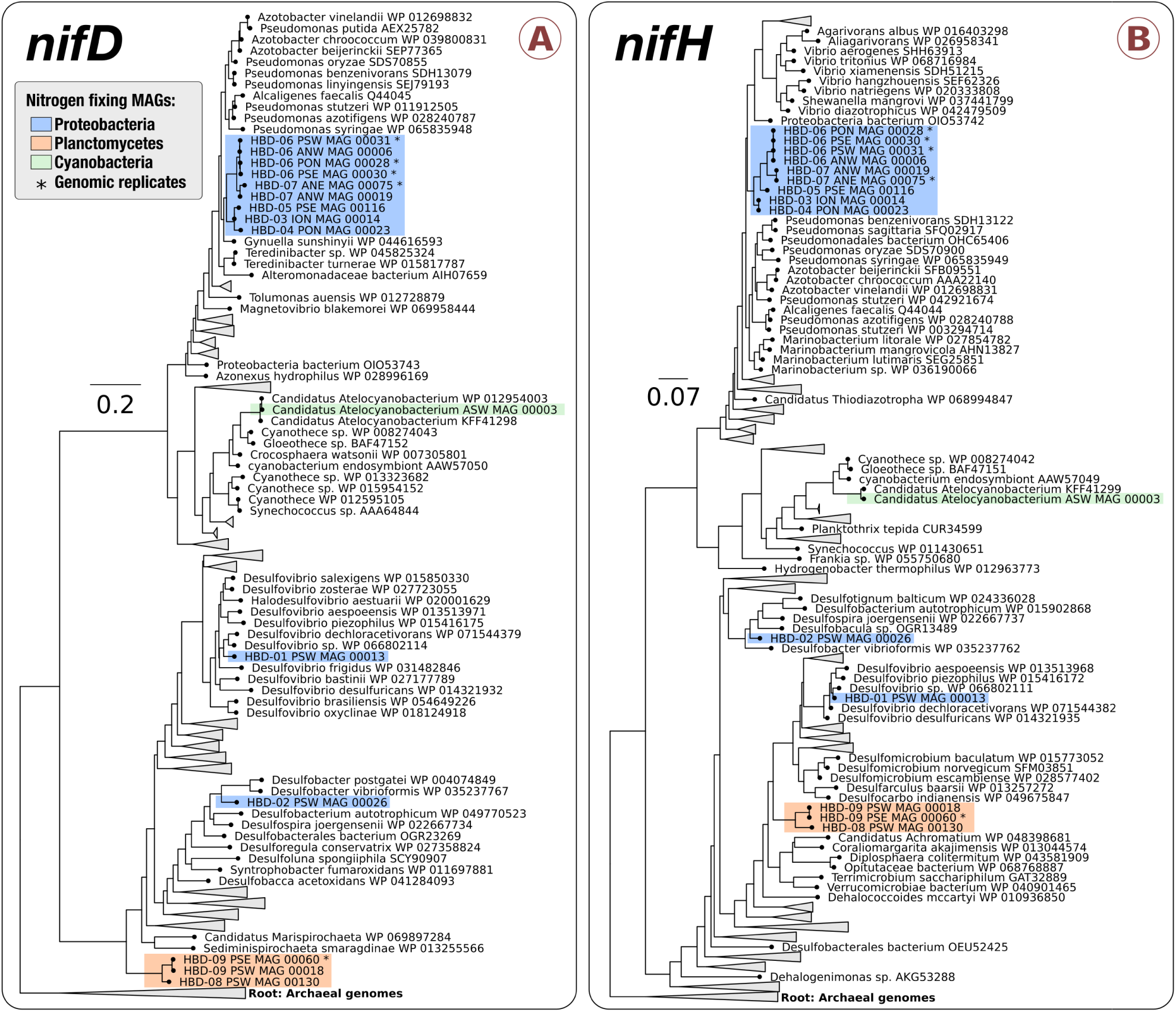
Phylogeny of nitrogen fixation genes. Phylogenetic analysis of *nifD* (A) and *nifH* (B) occurring in the 15 nitrogen fixing MAGs (including five redundant MAGs) we identified from TARA Oceans in relation to 252 and 316 reference genomes, respectively. MAGs are colored based on their phylogenetic affiliation at the phylum level.

### HBDs are not only diverse but also abundant in the surface ocean

The cumulative relative abundance of the Proteobacteria and Planctomycetes HBDs in the metagenomic dataset averaged to 0.01% and 0.05%, respectively (Table S3). In comparison, Ca. *A. thalassa* had a relative abundance averaging to 0.006%. The detection of Proteobacteria and Planctomycetes HBDs was very low in the Mediterranean Sea and Red Sea (0.00064% on average). In contrast, they were substantially enriched in metagenomes from the Pacific Ocean (0.14% in average) compared to the other regions (Figure 4). In fact, the Pacific Ocean metagenomes contained 81.4% of the 17.8 million reads that were recruited by HBD MAGs from the entire metagenomic dataset. In particular, the two most abundant Proteobacteria and Planctomycetes HBDs (HBD-06 and HBD-09) showed a broad distribution (Figure 4) and were significantly enriched in this ocean (Welch’s test; p-value *<*0.005). HBD-06 was also abundant in the northwest region of the Atlantic Ocean and to a lesser extent in the Southern Ocean, revealing that the ecological niche of a single HBD population can encompass multiple oceans and a wide range of temperatures (Table S3). Interestingly, HBD-07 and HBD-08, which are phylogenetically and functionally closely related to HBD-06 and HBD-09, respectively, were not only less abundant, but they also exhibited a different geographical distribution (Figure 4). We could not explain the increased signal for the nine HBDs in few geographic regions using temperature, salinity, or the concentration of essential inorganic chemicals including oxygen, phosphate and nitrate (Table S1).

**Figure 4.**
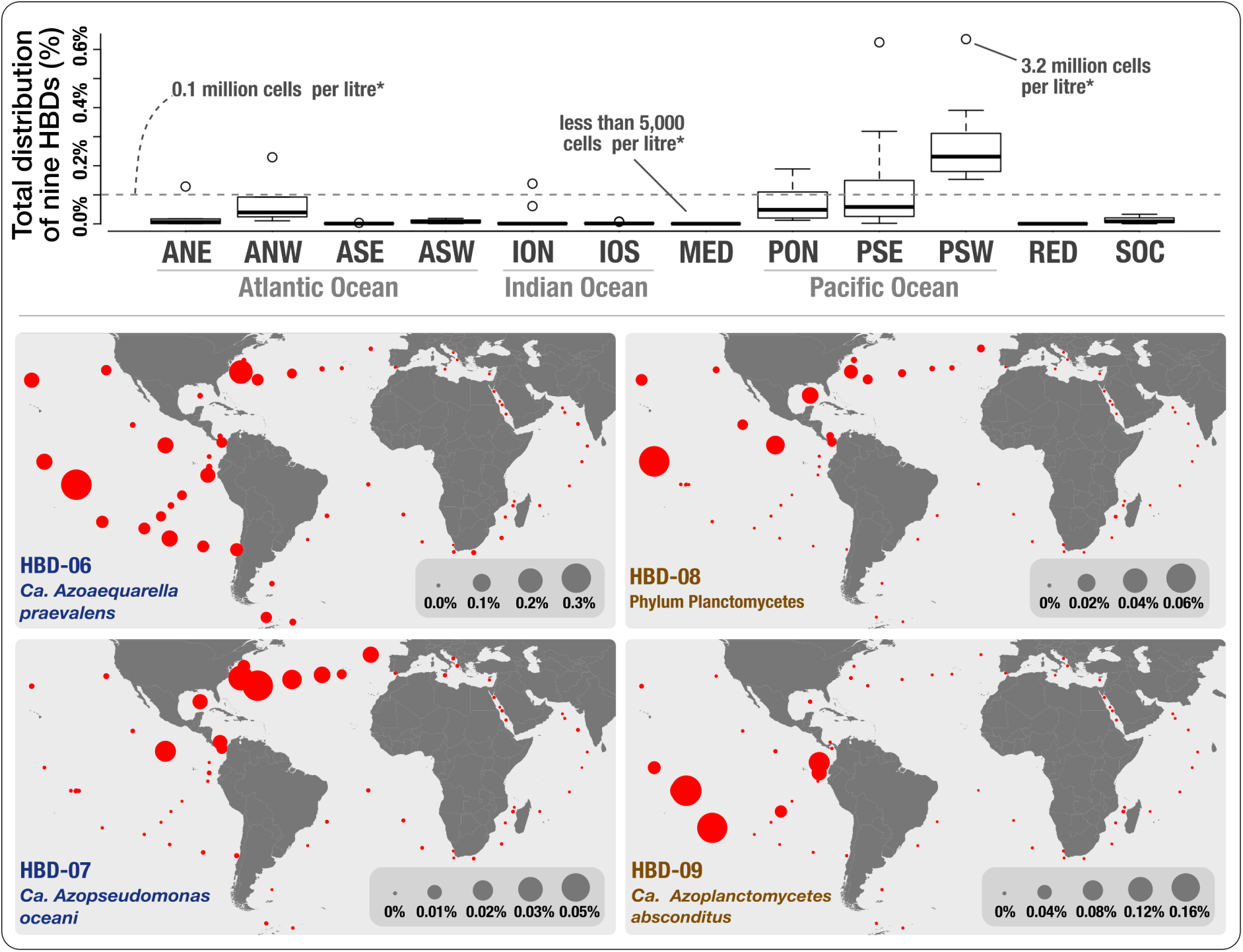
Nitrogen-fixing populations of Planctomycetes and Proteobacteria are abundant in the surface ocean. Cumulative relative distribution of the Planctomycetes (n=2) and Proteobacteria (n=7) HBDs in 93 metagenomes corresponding to 12 marine geographic regions (box-plots), and niche partitioning of HBD-06, HBD-07, HBD-08 and HBD-09 at the surface of four oceans and two seas (61 metagenomes from surface samples, world maps). *Assuming that each litre in the surface ocean contains 0.5 billion archaeal and bacterial cells [34].

To reconcile the abundance of nitrogen fixing populations in the surface of the open ocean with the inclusion of new HBDs described in this study, we used the previous PCR-based estimations of the abundance of non-cyanobacterial *nifH* genes. It is estimated that each litre of water in the surface ocean contains about 0.5 billion archaeal and bacterial cells [34]. Quantitative PCR surveys have estimated that all noncyanobacterial *nifH* genes combined generally range from 10 to 1,000 copies per litre and rarely reach 0.1 million copies [23, 24, 25, 26, 27]. One of the most abundant *nifH* phylotypes, gamma-24774A11, comprises up to 20,000 copies per litre in the Pacific Ocean, and is considered to represent one of the most important groups of non-cyanobacterial diazotrophs in the surface ocean [27, 35, 36]. In contrast, the same assumptions, and our abundance estimations based on metagenomic mapping, suggest that the nine populations of HBDs characterized in this study represent 0.5 million cells per litre on average (and up to 3.2 million cells) in the surface of the Pacific Ocean. HBD-06 alone represented on average more than 0.3 million cells per litre in this region. For a more direct comparison between relative abundance values derived from shotgun metagenomics and quantitative PCR surveys, we also recruited short metagenomic reads from TARA Oceans using a database of the *nifH* genes we identified in this study and gamma-24774A11 (Table S7). Mapping results showed that the *nifH* genes we identified in HBDs recruited 38.5 to 535 times more reads compared to gamma-24774A11 worldwide (i.e., when considering the 93 metagenomes altogether). These results altogether suggest that HBD populations can be up to about 2.5 orders of magnitude more abundant in the surface ocean than previously thought.

### A binomial naming for the HBDs we characterized from multiple geographic regions

Most of the MAGs we characterized in the present study correspond to novel genera, but the lack of culture representatives is preventing a formal taxonomical characterization of these new lineages. As an exception, we tentatively named the HBDs we independently characterized from multiple geographic regions (i.e., those for which we have genomic replicates) using the candidatus status and binomial naming system: *‘Candidatus Azoaequarella praevalens’* gen. nov., sp. nov., (HBD-06; representative genome is ANW 00006 with a completion of 98.1%) and *‘Ca. Azopseudomonas oceani’* gen. nov., sp. nov., (HBD-07; representative genome is ANW 00019 with a completion of 91.2%) within the order Pseudomonadales (unknown family), and *‘Ca. Azoplanctomyces absconditus’* gen. nov., sp. nov., (HBD-09; representative genome is PSW 00018 with a completion of 97.3%) within the Phylum Planctomycetes (unknown order and family). Note that ANI between ‘Ca. Azoaequarella praevalens’ and *‘Ca. Azopseudomonas oceani’* was 83.1%, supporting their assignment to different genera but suggesting they might belong to the same family. These three candidate species represent widespread and abundant nitrogen-fixing populations in the surface of the open ocean (Figure 4). Here is a brief explanation of their naming:

- *Azoaequarella praevalens* (N.L. n. azotum [from Fr. n. azote (from Gr. prep. a, not; Gr. n. zôê, life; N.Gr. n. azôê, not sustaining life)], nitrogen; N.L. pref. azo-, pertaining to nitrogen; L. v. aequare, to equalize; N.L. fem. n. Azoaequarella, the nitrogen equalizer; L. part. adj. praevalens, very powerful, very strong, here prevalent)
- *Azopseudomonas oceani* (N.L. n. azotum [from Fr. n. azote (from Gr. prep. a, not; Gr. n. zôê, life; N.Gr. n. azôê, not sustaining life)], nitrogen; N.L. pref. azo-, pertaining to nitrogen; Gr. adj. pseudês, false; Gr. fem. n. monas, a unit, monad; N.L. fem. n. Azopseudomonas, nitrogen-fixing false monad; L. gen. n. oceani, of the ocean)
- *Azoplanctomyces absconditus* (N.L. n. azotum [from Fr. n. azote (from Gr. prep. a, not; Gr. n. zôê, life; N.Gr. n. azôê, not sustaining life)], nitrogen; N.L. pref. azo-, pertaining to nitrogen; Gr. adj. planktos, wandering, floating; Gr. masc. n. mukês, fungus; N.L. masc. n. Azoplanctomyces, nitrogen-fixing floating fungus; L. part. adj. absconditus, hidden)

### Orphan *nifH* genes from other archaeal and bacterial phyla are also abundant

The non-redundant collection of 957 curated MAGs in which we searched for HBDs represent only 2.07% of the scaffolds in our assembly of TARA Oceans metagenomes. To identify more *nifH* genes, we also investigated the remaining ‘orphan’ scaffolds (42,193,607 genes). Our search based on amino acid similarity with the HBD database resulted in the recovery of nine additional non-redundant *nifH* genes (Table S7). Eight of them originated from the Pacific Ocean metagenomic co-assemblies, substantiating the unequal distribution patterns for nitrogen fixation genes we observed at the MAG level. The phylogenetic analysis on these *nifH* genes affiliated them with Elusimicrobia (n=2), Firmicutes (n=2), Proteobacteria (n=1), Spirochaeta (n=1), Verrucomicrobia (n=1), a group of uncultured bacteria (n=1), and Euryarchaeota (n=1) (Figure S1). This primer-independent analysis of *nifH* genes identified a small but phylogenetically remarkably wide range of lineages. In particular, the Euryarchaeota *nifH* (Id-1645572 from the region PON; 858 nucleotides-long) had several mismatches with commonly used *nifH* PCR primers [37]. Its best match in the NCBI non redundant database was a *nifH* gene of a Methanocalculus MAG (e-value=2e-93) obtained from an oil reservoir in Alaska [38]. These nine *nifH* genes had a mean coverage 21.7 to 56.3 times higher than gamma-24774A11 worldwide (Table S7). These findings suggest that although their genomic content has yet to be determined, abundant non-cyanobacterial diazotrophic populations in the surface ocean habitat are not limited to Planctomycetes and Proteobacteria.

## Discussion

The nine heterotrophic bacterial diazotrophs (HBDs) we describe in this study provide first genomic insights into widespread and abundant nitrogen-fixing surface ocean populations that are not affiliated with Cyanobacteria. These HBDs include two Planctomycetes populations, which represents the first observation of diazotrophs in this phylum. These findings complement decades of PCR surveys and cultivation efforts, and corroborate the relevance of metagenomic assembly and binning strategies to improve our understanding of microbial communities inhabiting the largest biome on Earth.

How relevant are these HBDs to the gross nitrogen budget of the world’s oceans? Although our study provides evidence to support the existence of thriving lineages of HBDs in the surface ocean, such as the recovery of near-identical population-level genomes from multiple geographic regions, it does not explain their role and impact on the nitrogen cycle. For instance, *‘Ca. Azoaequarella praevalens’* displayed functional capacity for both nitrogen fixation and denitrification, exposing a rare metabolic duality between nitrogen gain and loss. The remaining HBDs appear to lack genes for denitrification, however, the extent by which their relative abundances correlate with nitrogen gain for the surrounding microbial populations is unclear. These observations pose the necessity for more targeted analyses to understand the contribution of each HBD to nitrogen bioavailability in the surface ocean.

The HBDs we characterized were most abundant in the Pacific Ocean and the northwest of the Atlantic Ocean. One potential explanation for the differential distribution of HBDs at large scale could be a technical one: we may have failed to recover abundant nitrogen fixing populations from other geographic regions due to the limitations associated with metagenomic binning. To address this concern, we also focused on the occurrence of *nifH* genes in the assembled scaffolds that were not binned into any population genome and their abundance in the surface ocean. While this investigation revealed additional putative diazotrophs within other archaeal and bacterial phyla, their distribution patterns also mirrored those we observed at the level of population genomes. These findings altogether suggest that our study recovered the most abundant HBDs in the TARA Oceans metagenomes, and that their distribution may be driven by differential resource availability and niche specialization. Our analyses revealed a diverse group of HBDs enriched only in a few geographic regions. Although the available metadata did not seem to explain the niche boundaries for HBDs, the iron bioavailability may be playing a role: this element is particularly important for photosynthesis [39, 40, 41] and is known to represent a limiting factor for cyanobacterial diazotrophs in the Pacific Ocean [7]. Thus, marine systems co-limited by nitrogen and iron may represent appropriate ecological niches for HBDs, where they could be the main sources of nitrogen input into the surface ocean.

Overall, our study reveals that populations of HBDs within Planctomycetes and Proteobacteria, as well as putative diazotrophs within other archaeal and bacterial phyla, can be abundant in the surface ocean, occasionally across wide ecological niches spanning a large range of temperatures. Our investigation takes advantage of unprecedented amount of shotgun metagenomic sequencing data to investigate the diversity of *nifH* genes without primer bias. While our findings substantiate the previous observations made through PCR amplicon surveys regarding the diversity of HBDs in the surface of the oceans [23, 24, 25, 26, 27], they also demonstrate that amplicon surveys may have underestimated the abundance of HBDs by multiple orders of magnitude. This emphasizes the need to re-assess the role of heterotrophic bacteria in oceanic primary productivity and the nitrogen cycle. Cultivation efforts and additional environmental surveys will be essential to characterize the life-styles of these non-cyanobacterial populations, and to determine environmental conditions in which they fix nitrogen.

## Methods

The URL http://merenlab.org/data/2017 Delmont et al HBDs/ contains a reproducible work flow that extends the descriptions and parameters of programs used here for (1) metagenomic binning, (2) identification and curation of MAGs, (3) identification of Candidate Phyla Radiation MAGs, and (4) profiling of MAGs and *nifH* genes in the entire metagenomic dataset.

### TARA Oceans metagenomes

We acquired 93 metagenomes from the European Bioinformatics Institute (EBI) repository under the project ID ERP001736 and quality filtered the reads using the illumina-utils library [42] v1.4.1 (available from https://github.com/meren/illumina-utils). Noisy sequences were removed using the program ‘iu-filter-quality-minoche’ with default parameters, which implements a noise filtering described by Minoche et al. [43]. Table S1 reports accession numbers and additional information (including the number of reads and environmental metadata) for each metagenome.

### Metagenomic co-assemblies, gene calling, and binning

We organized the dataset into 12 ‘metagenomic sets’ based upon the geographic coordinates of metagenomes (Table S1). We co-assembled reads from each metagenomic set using MEGAHIT [44] v1.0.3, with a minimum scaffold length of 1 kbp, and simplified the scaffold header names in the resulting assembly outputs using anvi’o [33] v2.0.2 (available from http://merenlab.org/software/anvio). For each metagenomic set, we then binned scaffolds *>*2.5 kbp (*>*5 kbp for the Southern Ocean) following the workflow outlined in Eren et al.33. Briefly, (1) anvi’o profiled the scaffolds using Prodigal [45] v2.6.3 with default parameters to identify genes (Table S2), and HMMER [46] v3.1b2 to identify genes matching to archaeal [47] and bacterial [48, 49, 50, 51] single-copy core gene collections, (2) we used Centrifuge [52] with NCBI’s NT database to infer the taxonomy of genes (as described in http://merenlab.org/2016/06/18/importing-taxonomy), (3) we mapped short reads from the metagenomic set to the scaffolds using Bowtie2 [53] v2.0.5 and stored the recruited reads as BAM files using samtools [54], (4) anvi’o profiled each BAM file to estimate the coverage and detection statistics of each scaffold, and combined mapping profiles into a merged profile database for each metagenomic set. We then clustered scaffolds with the automatic binning algorithm CONCOCT [48] by constraining the number of clusters per metagenomic set to 100 to minimize the ‘fragmentation error’ (when multiple clusters describe one population), with the exception of the Southern Ocean (25 clusters) and the Pacific Ocean southeast (150 clusters) metagenomic sets. Finally, we manually binned each CONCOCT cluster (n=1,175) using the anvi’o interactive interface. See the Supplementary Information for details. Table S8 reports the genomic features (including completion and redundancy values) of the bins we characterized.

### Identification and curation of metagenome-assembled genomes (MAGs)

We defined all bins with *>*70% completion or *>*2 Mbp in length as MAGs (Table S2). We then individually refined each MAG as outlined in Delmont and Eren [55], and renamed scaffolds they contained accordingly to their MAG ID in order to insure that the names of all scaffolds in MAGs we characterized from the 12 metagenomic sets were unique.

### Taxonomic and functional inference of MAGs

We used CheckM [56] to infer the taxonomy of MAGs based on the proximity of 43 single-copy gene markers within a reference genomic tree. We also used Centrifuge, RAST [57] and manual BLAST searches of single copy core genes against the NCBI’s nonredundant database to manually refine the CheckM taxonomic inferences, especially regarding the archaeal and eukaryotic MAGs. We also used the occurrence of bacterial single-copy genes to identify MAGs affiliated to the Candidate Phyla Radiation (as described in http://merenlab.org/2016/04/17/predicting-CPRGenomes/). Table S4 reports our curated taxonomic inference of MAGs. We used KEGG (the 04/14/2014 release) to identify functions and pathways in MAGs. We also used RAST to identify functions in 15 MAGs that contained the complete set of nitrogen fixation genes (originally identified from the KEGG pathways). Tables S6 and S9 report the RAST and KEEG results, respectively. We used Gephi [58] v0.8.2 to generate a functional network using the Force Atlas 2 algorithm to connect MAGs and RAST functions. We correlated node sizes to the number of edges they contained, which resulted in larger nodes for MAGs compared to functions.

### Characterization of a non-redundant database of MAGs

We concatenated all scaffolds from the genomic database of MAGs into a single FASTA file and used Bowtie2 and samtools to recruit and store reads from the 93 metagenomes. We used anvi’o to determine the coverage values, detection and relative distribution of MAGs and individual genes across metagenomes (Table S10). We determined the Pearson correlation coefficient of each pair of MAGs based upon their relative distribution across the 93 metagenomes using the function ‘cor’ in R [59] (Table 5). We finally used NUCmer [60] to determine the average nucleotide identity (ANI) of each pair of MAG affiliated to the same phylum for improved performance (the Proteobacteria MAGs were further split at the class level) (Table 5). MAGs were considered redundant when their ANI reached to 99% (minimum alignment of *>*75% of the smaller genome in each comparison) and the Pearson correlation coefficient above 0.9. We then selected a single MAG to represent a group of redundant MAGs based on the largest ‘completion minus redundancy’ value from single-copy core genes for Archaea and Bacteria, or longer genomic length for Eukaryota. This analysis provided a non-redundant genomic database of MAGs. We performed a final mapping of all metagenomes to calculate the mean coverage and detection of these MAGs (Table S3, Supplementary Information).

### Statistical analyses

We used STAMP [61] and Welch’s test to identify non-redundant MAGs that were significantly enriched in the Pacific Ocean compared to all the other regions combined. Table S3 reports the p-values for each MAG.

### World maps

We used the ggplot2 [62] package for R to visualize the metagenomic sets and relative distribution of MAGs in the world map.

### Phylogenomic analysis of MAGs

We used PhyloSift [63] v1.0.1 with default parameters to infer associations between MAGs in a phylogenomic context. Briefly, PhyloSift (1) identifies a set of 37 marker gene families in each genome, (2) concatenates the alignment of each marker gene family across genomes, and (3) computes a phylogenomic tree from the concatenated alignment using FastTree [64] v2.1. We rooted the phylogenomic tree to the phylum Planctomycetes with FigTree [65] v1.4.3, and used anvi’o to visualize it with additional data layers.

### Identification of additional *nifH* sequences in orphan scaffolds

We used DIAMOND [66] to generate a database of *nifH* genes we identified in the nine HBDs, and to search for additional *nifH* amino acid sequences within the genes Prodigal identified in scaffolds longer than 1,000 nucleotides. We considered only hits with an e-value of *<*1e-50, and defined them as *nifH* genes only when (1) ‘nitrogenase’ was the top blastx hit against the NCBI’s nr database, and (2) the characteristic [4Fe-4S]-binding site (Prosite signature PDOC00580) was present in their amino acid sequence.

### Mean coverage of *nifH* genes across 93 metagenomes

We concatenated all *nifH* genes (orphan genes as well as the ones affiliated with HBDs) into a single FASTA file along with the additional reference sequence gamma-24774A11. To estimate the mean coverage of all *nifH* sequences, we recruited reads from all metagenomes and profiled the resulting mapping results with anvi’o as described above. Table S7 reports the mean coverage estimates.

### Phylogenetic analysis of *nifD* and *nifH* genes

We imported the amino acid sequences of *nifD* and *nifH* genes identified in this study into ARB v.5.5-org-9167 [67], which built databases by identifying representative, non-redundant protein reference sequences by blastp searches against the NCBI nr database. In ARB, we aligned sequences to each other using ClustalW [68], manually refined alignments, and calculated phylogenetic trees with PhyML [69] using the ‘WAG’ amino acid substitution model, and a 10% conservation filter.

### Figure designs

We used the open-source vector graphics editor Inskape (available from http://inkscape.org/) to finalize all the figures.

### Data availability

We stored scaffolds *>*2.5 kbp we generated from the 12 metagenomic co-assemblies in the NCBI Bioproject PRJNA326480. We also made publicly available the raw assembly results that include scaffolds *>*1 kbp (doi:10.6084/m9.figshare.4902920), amino acid sequences for 42.2 million genes identified in raw assembly results (doi:10.6084/m9.figshare.4902917), the FASTA files for our final collection of 957 non-redundant MAGs (doi:10.6084/m9.figshare.4902923), the extensive anvi’o summary output for non-redundant MAGs and their distribution across metagenomes (doi:10.6084/m9.figshare.4902926), and finally the self-contained anvi’o split profiles for each non-redundant MAG (doi:10.6084/m9.figshare.4902941).

All supplementary information is available via doi:10.6084/m9.figshare.4902938

## Author’s contributions

T.O.D. and A.M.E. conceived the study, developed the analysis tools, and wrote the paper. All authors contributed to the analysis of the data, and reviewed drafts of the paper.

## Acknowledgements

We thank the TARA Oceans consortium for generating metagenomic datasets of great legacy. We thank Mike Lee for helpful discussions, and our computer systems administrator Rich Fox for his patience and help. This study was supported by the Frank R. Lillie Research Innovation Award and the startup funds from the University of Chicago to AME, and a Netherlands Organization for Scientific Research grant (VENI 863.14.019) to SL.

## References

1. Charlson, R.J., Lovelock, J.E., Andreae, M.O., Warren, S.G.: Oceanic phytoplankton, atmospheric sulphur, cloud albedo and climate. Nature 326(|mApril 1987), 655–661 (1987). doi:10.1038/326655a0

2. Falkowski, P.G., Barber, R.T., Smetacek, V.: Biogeochemical controls and feedbacks on ocean primary production. Science 281(1998), 200–206 (1998). doi:10.1126/science.281.5374.200

3. Arrigo, K.R.: Marine microorganisms and global nutrient cycles. Nature 437(7057), 349–355 (2005)

4. Moore, C.M., Mills, M.M., Arrigo, K.R., Berman-Frank, I., Bopp, L., Boyd, P.W., Galbraith, E.D., Geider, R.J., Guieu, C., Jaccard, S.L., Jickells, T.D., La Roche, J., Lenton, T.M., Mahowald, N.M., Maranon, E., Marinov, I., Moore, J.K., Nakatsuka, T., Oschlies, A., Saito, M.A., Thingstad, T.F., Tsuda, A., Ulloa, O.: Processes and patterns of oceanic nutrient limitation. Nature Geosci 6(9), 701–710 (2013). doi:10.1038/ngeo1765

5. Tyrrell, T.: The relative influences of nitrogen and phosohorus on oceanic primary production. Nature 400(6744), 525–531 (1999). doi:10.1038/22941

6. Capone, D.G.: Trichodesmium, a Globally Significant Marine Cyanobacterium. Science 276(5316), 1221–1229 (1997). doi:10.1126/science.276.5316.1221

7. Sohm, J.a., Webb, E.a., Capone, D.G.: Emerging patterns of marine nitrogen fixation. Nature reviews. Microbiology 9(7), 499–508 (2011). doi:10.1038/nrmicro2594

8. Dos Santos, P.C., Fang, Z., Mason, S.W., Setubal, J.C., Dixon, R.: Distribution of nitrogen fixation and nitrogenase-like sequences amongst microbial genomes. BMC Genomics 13(1), 162 (2012). doi:10.1186/1471-2164-13-162

9. Zehr, J.P., Jenkins, B.D., Short, S.M., Steward, G.F.: Nitrogenase gene diversity and microbial community structure: a cross-system comparison. Environ Microbiol 5(7), 539–554 (2003). doi:10.1046/j.1462-2920.2003.00451.x

10. Karl, D., Letelier, R., Tupas, L., Dore, J., Christian, J., Hebel, D.: The role of nitrogen fixation in biogeochemical cycling in the subtropical North Pacific Ocean. Nature 388(6642), 533–538 (1997). doi:10.1038/41474

11. Carpenter, E.J., Romans, K.: Major role of the cyanobacterium trichodesmium in nutrient cycling in the north atlantic ocean. Science (New York, N.Y.) 254(1989), 1356–1358 (1991). doi:10.1126/science.254.5036.1356

12. Carpenter, E.J., Capone, D.G.: Nitrogen fixation in Trichodesmium blooms. In: Marine Pelagic Cyanobacteria: Trichodesmium and Other Diazotrophs, pp. 211–217 (1992). doi:10.1007/978-94-015-7977-313

13. Carpenter, E.J., Price, C.C.: Nitrogen fixation, distribution, and production of *Oscillatoria (Trichodesmium)* spp. in the western Sargasso and Caribbean Seas. Limnology and Oceanography 22(1), 60–72 (1977). doi:10.4319/lo.1977.22.1.0060

14. Zehr, J.P., Waterbury, J.B., Turner, P.J., Montoya, J.P., Omoregie, E., Steward, G.F., Hansen, A., Karl, D.M.: Unicellular cyanobacteria fix N2 in the subtropical North Pacific Ocean. Nature 412(6847), 635–638 (2001). doi:10.1038/35088063

15. Lehtimäki, J., Moisander, P., Sivonen, K., Kononen, K.: Growth, nitrogen fixation, and Nodularin production by two Baltic Sea cyanobacteria. Applied and Environmental Microbiology 63(5), |p1647–1656 (1997)

16. Wyatt, J.T., Silvey, J.K.: Nitrogen fixation by gloeocapsa. Science (New York, N.Y.) 165(3896), 908–9 (1969). doi:10.1126/science.165.3896.908

17. Carpenter, E.J., Capone, D.G., Rueter, J.G.: Marine Pelagic Cyanobacteria: Trichodesmium and Other Diazotrophs., p. 358 (1992). doi:10.1007/978-94-015-7977-3. arXiv:1011.1669v3. https://books.google.co.uk/books?hl=en&lr=&id=CXnoCAAAQBAJ&oi=fnd&pg=PA1&dq=Marine+Pelagic+Cyanobacteria:+Trichodesmium+and+other+Diazo

18. Kaneko, T., Nakamura, Y., Wolk, C.P., Kuritz, T., Sasamoto, S., Watanabe, a., Iriguchi, M., Ishikawa, a., Kawashima, K., Kimura, T., Kishida, Y., Kohara, M., Matsumoto, M., Matsuno, a., Muraki, a., Nakazaki, N., Shimpo, S., Sugimoto, M., Takazawa, M., Yamada, M., Yasuda, M., Tabata, S.: Complete genomic sequence of the filamentous nitrogen-fixing cyanobacterium Anabaena sp. strain PCC 7120. DNA research : an international journal for rapid publication of reports on genes and genomes 8(5), 205–213227253 (2001). doi:10.1093/dnares/8.5.205

19. Tripp, H.J., Bench, S.R., Turk, K.a., Foster, R.a., Desany, B.a., Niazi, F., Affourtit, J.P., Zehr, J.P.: Metabolic streamlining in an open-ocean nitrogen-fixing cyanobacterium. Nature 464(7285), 90–94 (2010). doi:10.1038/nature08786

20. Dyhrman, S.T., Chappell, P.D., Haley, S.T., Moffett, J.W., Orchard, E.D., Waterbury, J.B., Webb, E.A.: Phosphonate utilization by the globally important marine diazotroph *Trichodesmium*. Nature 439(7072), 68–71 (2006). doi:10.1038/nature04203

21. Zehr, J.P., Mellon, M.T., Zani, S.: New nitrogen-fixing microorganisms detected in oligotrophic oceans by amplification of nitrogenase (nifH) genes. Applied and Environmental Microbiology 64(9), 3444–3450 (1998). doi:PMC106745

22. Riemann, L., Farnelid, H., Steward, G.F.: Nitrogenase genes in non-cyanobacterial plankton: Prevalence, diversity and regulation in marine waters (2010). doi:10.3354/ame01431

23. Church, M.J., Short, C.M., Jenkins, B.D., Karl, D.M., Zehr, J.P.: Temporal patterns of nitrogenase gene (nifH) expression in the oligotrophic North Pacific Ocean. Applied and Environmental Microbiology 71(9), 5362–5370 (2005). doi:10.1128/AEM.71.9.5362-5370.2005

24. Church, M.J., Björkman, K.M., Karl, D.M., Saito, M.a., Zehr, J.P.: Regional distributions of nitrogen-fixing bacteria in the Pacific Ocean. Limnology and Oceanography 53(1), 63–77 (2008). doi:10.4319/lo.2008.53.1.0063

25. Zehr, J.P., Montoya, J.P., Jenkins, B.D., Hewson, I., Mondragon, E., Short, C.M., Church, M.J., Hansen, A., Karl, D.M.: Experiments linking nitrogenase gene expression to nitrogen fixation in the North Pacific subtropical gyre

26. Fong, A.A., Karl, D.M., Lukas, R., Letelier, R.M., Zehr, J.P., Church, M.J.: Nitrogen fixation in an anticyclonic eddy in the oligotrophic North Pacific Ocean. The ISME journal 2(6), 663–676 (2008). doi:10.1038/ismej.2008.22

27. Moisander, P.H., Beinart, R.A., Voss, M., Zehr, J.P.: Diversity and abundance of diazotrophic microorganisms in the South China Sea during intermonsoon. The ISME Journal 251, 954–967 (2008). doi:10.1038/ismej.2008.51

28. Bombar, D., Paerl, R.W., Riemann, L.: Marine Non-Cyanobacterial Diazotrophs: Moving beyond Molecular Detection (2016). doi:10.1016/j.tim.2016.07.002

29. Sunagawa, S., Coelho, L.P., Chaffron, S., Kultima, J.R., Labadie, K., Salazar, G., Djahanschiri, B., Zeller, G., Mende, D.R., Alberti, A., Cornejo-Castillo, F.M., Costea, P.I., Cruaud, C., D’Ovidio, F., Engelen, S., Ferrera, I., Gasol, J.M., Guidi, L., Hildebrand, F., Kokoszka, F., Lepoivre, C., Lima-Mendez, G., Poulain, J., Poulos, B.T., Royo-Llonch, M., Sarmento, H., Vieira-Silva, S., Dimier, C., Picheral, M., Searson, S., Kandels-Lewis, S., Bowler, C., de Vargas, C., Gorsky, G., Grimsley, N., Hingamp, P., Iudicone, D., Jaillon, O., Not, F., Ogata, H., Pesant, S., Speich, S., Stemmann, L., Sullivan, M.B., Weissenbach, J., Wincker, P., Karsenti, E., Raes, J., Acinas, S.G., Bork, P.: Ocean plankton. Structure and function of the global ocean microbiome. Science (New York, N.Y.) 348(6237), 1261359 (2015). doi:10.1126/science.1261359

30. Zehr, J.P., Bench, S.R., Carter, B.J., Hewson, I., Niazi, F., Shi, T., Tripp, H.J., Affourtit, J.P.: Globally Distributed Uncultivated Oceanic N2-Fixing Cyanobacteria Lack Oxygenic Photosystem II. Science 322(5904), 1110–1112 (2008). doi:10.1126/science.1165340

31. Thompson, A.W., Foster, R.A., Krupke, A., Carter, B.J., Musat, N., Vaulot, D., Kuypers, M.M.M., Zehr, J.P.: Unicellular cyanobacterium symbiotic with a single-celled eukaryotic alga. Science (New York, N.Y.) 337(6101), 1546–50 (2012). doi:10.1126/science.1222700

32. Delmont, T.O., Eren, A.M., Maccario, L., Prestat, E., Esen, Ö.C., Pelletier, E., Le Paslier, D., Simonet, P., Vogel, T.M.: Reconstructing rare soil microbial genomes using in situ enrichments and metagenomics. Frontiers in microbiology 6, 358 (2015). doi:10.3389/fmicb.2015.00358

33. Eren, A.M., Esen, Ö.C., Quince, C., Vineis, J.H., Morrison, H.G., Sogin, M.L., Delmont, T.O.: Anvi’o: an advanced analysis and visualization platform for ‘omics data. PeerJ 3, 1319 (2015). doi:10.7717/peerj.1319

34. Whitman, W.B., Coleman, D.C., Wiebe, W.J.: Prokaryotes: the unseen majority. Proceedings of the National Academy of Sciences of the United States of America 95(12), 6578–6583 (1998)

35. Moisander, P.H., Serros, T., Paerl, R.W., Beinart, R.a., Zehr, J.P.: Gammaproteobacterial diazotrophs and nifH gene expression in surface waters of the South Pacific Ocean. The ISME journal 8(10), 1962–1973 (2014). doi:10.1038/ismej.2014.49

36. Langlois, R., Großkopf, T., Mills, M., Takeda, S., LaRoche, J.: Widespread distribution and expression of Gamma A (UMB), an uncultured, diazotrophic, ??-proteobacterial nifH phylotype. PLoS ONE 10(6) (2015). doi:10.1371/journal.pone.0128912

37. Zehr, J.P., Turner, P.J.: Nitrogen Fixation : Nitrogenase Genes and Gene Expression. METHODS IN MICROBIOLOGY, Volume 30, 271–286 (2001)

38. Hu, P., Tom, L., Singh, A., Thomas, B.C., Baker, B.J., Piceno, Y.M., Andersen, G.L., Banfield, J.F.: Genome-resolved metagenomic analysis reveals roles for candidate phyla and other microbial community members in biogeochemical transformations in oil reservoirs. mBio 7(1) (2016). doi:10.1128/mBio.01669-15

39. Ferreira, F., Straus, N.A.: Iron deprivation in cyanobacteria. Journal of Applied Phycology 6(2), 199–210 (1994). doi:10.1007/BF02186073

40. Basu, P., Stolz, J., Croot, P., Sunda, W.G.: High iron requirement for growth, photosynthesis, and low-light acclimation in the coastal cyanobacterium Synechococcus bacillaris (2015). doi:10.3389/fmicb.2015.00561

41. Mills, M.M., Ridame, C., Davey, M., La Roche, J., Geider, R.J.: Iron and phosphorus co-limit nitrogen fixation in the eastern tropical North Atlantic. Nature 429(6989), 292–294 (2004). doi:10.1038/nature02550

42. Eren, A.M., Vineis, J.H., Morrison, H.G., Sogin, M.L.: A Filtering Method to Generate High Quality Short Reads Using Illumina Paired-End Technology. PLoS ONE 8(6), 66643 (2013). doi:10.1371/journal.pone.0066643

43. Minoche, A.E., Dohm, J.C., Himmelbauer, H.: Evaluation of genomic high-throughput sequencing data generated on Illumina HiSeq and genome analyzer systems. Genome biology 12(11), 112 (2011). doi:10.1186/gb-2011-12-11-r112

44. Li, D., Liu, C.M., Luo, R., Sadakane, K., Lam, T.W.: MEGAHIT: An ultra-fast single-node solution for large and complex metagenomics assembly via succinct de Bruijn graph. Bioinformatics 31(10), 1674–1676 (2014). doi:10.1093/bioinformatics/btv033. 1401.7457

45. Hyatt, D., Chen, G.-L., Locascio, P.F., Land, M.L., Larimer, F.W., Hauser, L.J.: Prodigal: prokaryotic gene recognition and translation initiation site identification. BMC bioinformatics 11, 119 (2010). doi:10.1186/1471-2105-11-119

46. Eddy, S.R.: Accelerated Profile HMM Searches. PLoS computational biology 7(10), 1002195 (2011). doi:10.1371/journal.pcbi.1002195

47. Rinke, C., Schwientek, P., Sczyrba, A., Ivanova, N.N., Anderson, I.J., Cheng, J.-F., Darling, A.E., Malfatti, S., Swan, B.K., Gies, E.a., Dodsworth, J.a., Hedlund, B.P., Tsiamis, G., Sievert, S.M., Liu, W.-T., Eisen, J.a., Hallam, S.J., Kyrpides, N.C., Stepanauskas, R., Rubin, E.M., Hugenholtz, P., Woyke, T.: Insights into the phylogeny and coding potential of microbial dark matter. Nature 499(7459), 431–437 (2013). doi:10.1038/nature12352.3313

48. Alneberg, J., Bjarnason, B.S., de Bruijn, I., Schirmer, M., Quick, J., Ijaz, U.Z., Lahti, L., Loman, N.J., Andersson, A.F., Quince, C.: Binning metagenomic contigs by coverage and composition. Nature Methods 11(11), 1144–1146 (2014). doi:10.1038/nmeth.3103

49. Campbell, J.H., O’Donoghue, P., Campbell, A.G., Schwientek, P., Sczyrba, A., Woyke, T., Söll, D., Podar, M.: UGA is an additional glycine codon in uncultured SR1 bacteria from the human microbiota. Proceedings of the National Academy of Sciences of the United States of America 110(14), 5540–5545 (2013). doi:10.1073/pnas.1303090110

50. Dupont, C.L., Rusch, D.B., Yooseph, S., Lombardo, M.-J., Richter, R.A., Valas, R., Novotny, M., Yee-Greenbaum, J., Selengut, J.D., Haft, D.H., Halpern, A.L., Lasken, R.S., Nealson, K., Friedman, R., Venter, J.C.: Genomic insights to SAR86, an abundant and uncultivated marine bacterial lineage. The ISME journal 6(6), 1186–1199 (2012). doi:10.1038/ismej.2011.189

51. Creevey, C.J., Doerks, T., Fitzpatrick, D.A., Raes, J., Bork, P.: Universally distributed single-copy genes indicate a constant rate of horizontal transfer. PLoS one 6(8), 22099 (2011). doi:10.1371/journal.pone.0022099

52. Kim, D., Song, L., Breitwieser, F.P., Salzberg, S.L.: Centrifuge: rapid and sensitive classification of metagenomic sequences. bioRxiv, 054965 (2016). doi:10.1101/054965

53. Langmead, B., Salzberg, S.L.: Fast gapped-read alignment with Bowtie 2. Nature methods 9(4), 357–359 (2012). doi:10.1038/nmeth.1923

54. Li, H., Handsaker, B., Wysoker, A., Fennell, T., Ruan, J., Homer, N., Marth, G., Abecasis, G., Durbin, R.: The Sequence Alignment/Map format and SAMtools. Bioinformatics (Oxford, England) 25(16), 2078–2079 (2009). doi:10.1093/bioinformatics/btp352

55. Delmont, T.O., Eren, A.M.: Identifying contamination with advanced visualization and analysis practices: metagenomic approaches for eukaryotic genome assemblies. PeerJ 4, 1839 (2016). doi:10.7717/peerj.1839

56. Parks, D.H., Imelfort, M., Skennerton, C.T., Hugenholtz, P., Tyson, G.W.: CheckM: assessing the quality of microbial genomes recovered from isolates, single cells, and metagenomes. Genome research 25(7), 1043–1055 (2015). doi:10.1101/gr.186072.114

57. Aziz, R.K., Bartels, D., Best, A.A., DeJongh, M., Disz, T., Edwards, R.A., Formsma, K., Gerdes, S., Glass, E.M., Kubal, M., Meyer, F., Olsen, G.J., Olson, R., Osterman, A.L., Overbeek, R.A., McNeil, L.K., Paarmann, D., Paczian, T., Parrello, B., Pusch, G.D., Reich, C., Stevens, R., Vassieva, O., Vonstein, V., Wilke, A., Zagnitko, O.: The RAST Server: rapid annotations using subsystems technology. BMC genomics 9, 75 (2008). doi:10.1186/1471-2164-9-75

58. Bastian, M., Heymann, S., Jacomy, M.: Gephi: an open source software for exploring and manipulating networks. ICWSM 2, 361–362 (2009). doi:10.1136/qshc.2004.010033

59. R Development Core Team, R.: R: A Language and Environment for Statistical Computing (2011). doi:10.1007/978-3-540-74686-7. http://www.r-project.org

60. Delcher, A.L., Phillippy, A., Carlton, J., Salzberg, S.L.: Fast algorithms for large-scale genome alignment and comparison. Nucleic acids research 30(11), 2478–2483 (2002). doi:10.1093/nar/30.11.2478

61. Parks, D.H., Beiko, R.G.: Identifying biologically relevant differences between metagenomic communities. Bioinformatics (Oxford, England) 26(6), 715–721 (2010). doi:10.1093/bioinformatics/btq041

62. Ginestet, C.: ggplot2: Elegant Graphics for Data Analysis. Journal of the Royal Statistical Society: Series A (Statistics in Society) 174(1), 245–246 (2011). doi:10.1111/j.1467-985X.2010.006769.x

63. Darling, A.E., Jospin, G., Lowe, E., Matsen, F.A., Bik, H.M., Eisen, J.A.: PhyloSift: phylogenetic analysis of genomes and metagenomes. PeerJ 2, 243 (2014). doi:10.7717/peerj.243

64. Price, M.N., Dehal, P.S., Arkin, A.P.: FastTree 2 - Approximately maximum-likelihood trees for large alignments. PLoS ONE 5(3) (2010). doi:10.1371/journal.pone.0009490

65. Rambaut, A.: FigTree, a graphical viewer of phylogenetic trees. Institute of Evolutionary Biology University of Edinburgh (2009)

66. Buchfink, B., Xie, C., Huson, D.H.: Fast and sensitive protein alignment using DIAMOND. Nature methods 12(1), 59–60 (2015). doi:10.1038/nmeth.3176

67. Ludwig, W., Strunk, O., Westram, R., Richter, L., Meier, H., Yadhukumar, A., Buchner, A., Lai, T., Steppi, S., Jacob, G., Förster, W., Brettske, I., Gerber, S., Ginhart, A.W., Gross, O., Grumann, S., Hermann, S., Jost, R., König, A., Liss, T., Lüßbmann, R., May, M., Nonhoff, B., Reichel, B., Strehlow, R., Stamatakis, A., Stuckmann, N., Vilbig, A., Lenke, M., Ludwig, T., Bode, A., Schleifer, K.H.: ARB: A software environment for sequence data. Nucleic Acids Research 32(4), 1363–1371 (2004). doi:10.1093/nar/gkh293

68. Larkin, M.A., Blackshields, G., Brown, N.P., Chenna, R., Mcgettigan, P.A., McWilliam, H., Valentin, F., Wallace, I.M., Wilm, A., Lopez, R., Thompson, J.D., Gibson, T.J., Higgins, D.G.: Clustal W and Clustal X version 2.0. Bioinformatics 23(21), 2947–2948 (2007). doi:10.1093/bioinformatics/btm404.1011.1669

69. Guindon, S., Dufayard, J.F., Lefort, V., Anisimova, M., Hordijk, W., Gascuel, O.: New algorithms and methods to estimate maximum-likelihood phylogenies: Assessing the performance of PhyML 3.0. Systematic Biology 59(3), 307–321 (2010). doi:10.1093/sysbio/syq010

